# RNA secondary structure prediction with Convolutional Neural Networks

**DOI:** 10.1101/2021.05.24.445408

**Authors:** Mehdi Saman Booy, Alexander Ilin, Pekka Orponen

## Abstract

Predicting the secondary, i.e. base-pairing structure of a folded RNA strand is an important problem in synthetic and computational biology. First-principle algorithmic approaches to this task are challenging because existing models of the folding process are inaccurate, and even if a perfect model existed, finding an optimal solution would be in general NP-complete. In this paper, we propose a simple, yet extremely effective data-driven approach. We represent RNA sequences in the form of three-dimensional tensors in which we encode possible relations between all pairs of bases in a given sequence. We then use a convolutional neural network to predict a two-dimensional map which represents the correct pairings between the bases. Our model achieves significant accuracy improvements over existing methods on two standard datasets. Our experiments show excellent performance of the model across a wide range of sequence lengths and RNA families. We also observe considerable improvements in predicting complex pseudoknotted RNA structures, as compared to previous approaches.

**Author summary:** Structure prediction for RNA sequences is a computationally difficult task that is of increasing importance in applications such as medical diagnostics and drug design; this is because the structure of a folded RNA strand to a large extent defines its function. An open RNA strand can fold to many different structures of varying thermal stability, and the goal of structure prediction is to determine a most stable one among these. There are two main difficulties to this task. Firstly, a given RNA sequence can fold into an enormous number of alternative structures, and a computational search for a most stable one in this huge space can be very demanding. The search can however be facilitated by using heuristics that take into account some underlying principles of the folding process. Here is where machine learning methods come into play: they are suitable for discovering patterns in data, and can thus predict features of the desired structure based on previously learned patterns. Secondly, there do not yet exist fully satisfactory coarse-grained models for the most popular metric for stability, the free energy of the folded structure. Although in principle a minimum free energy (MFE) structure should be a good candidate for a most stable one, MFE structures determined according to current energy models do not match experimental data on native RNA conformations very well. We show how to use an artificial neural network design to predict the structure for a given RNA sequence with high accuracy only by learning from samples whose native structures have been experimentally characterized, independent of any stability metric or energy model.

## Introduction

### The RNA structure prediction problem

RNA is a highly versatile molecule of life: it has several key roles in the essential cellular processes of gene expression and regulation, carries cellular signals, and serves as a multi-purpose catalyst. It is a linear polymeric molecule constituted of elementary nucleotide units with bases adenine (A), cytosine (C), guanine (G) and uracil (U), bound to a sugar-phosphate backbone. RNA molecules, which are natively single-stranded, fold upon themselves to create biologically active 3D conformations, following mostly (but not completely) similar Watson-Crick base pairing rules as DNA: adenine pairs with uracil (A-U) and guanine with cytosine (G-C), but often also with uracil (the G-U *wobble pair*). To understand, and eventually control, this critical function-forming process, it is important to be able to predict how a given nucleotide sequence (the *primary structure*) folds upon itself to create a base-pairing *secondary structure* and eventually the geometric 3D *tertiary structure*. Because predicting the final tertiary structure is extraordinarily difficult, much research has focused on trying to resolve the intermediate problem of secondary structure formation.

In simple cases, RNA secondary structures exhibit a clean hierarchical arrangement composed of blocks of matching base-pairs (*stem segments*) interspersed with intervals of unpaired bases (*loops*), analogous to a well-parenthesised string in a formal language. In fact, one standard representation for these basic structures is the *dot-bracket notation*, where the bases are enumerated from the 5’-sugar end of the backbone towards the 3’-sugar end: each base initiating a pair is denoted by an opening parenthesis, the matching closing base by a closing parenthesis, and the unpaired bases by dots. The situation is, however, significantly complicated by base-pairs that break this hierarchical arrangement, so called *pseudoknot* connections. Theoretically, an optimal non-pseudoknotted secondary structure for a given sequence can be found efficiently by a dynamic programming approach, whereas the problem becomes NP-complete when pseudoknots are allowed.

### Related work

Most secondary structure prediction approaches propose some scoring function and strive to find appropriate structures with respect to this function. In the common case where the score is based on an energy model, the goal is either to determine a minimum free energy (MFE) structure in the given model, or sample structures according to a corresponding Boltzmann probability distribution.

#### Energy-based algorithmic methods

These methods find a thermodynamically minimum free energy structure for a given sequence and an energy model. Zuker [1, 2] proposed a basic dynamic programming approach to find an MFE structure by aggregating locally optimal structural elements with respect to a proposed energy model. Later on, Turner [3, 4] presented a more comprehensive “nearest neighbour” energy model, which became the core for many other methods originating from the Zuker algorithm, such as UNAFold [5], RNAStructure [6] and Vienna RNAfold [7], the latter tuning the energy parameters somewhat. Lyngsø and Pedersen [8] showed that finding MFE structures in a given energy model becomes NP-complete when pseudoknots are allowed. Hence, algorithmic methods based on dynamic programming cannot cover pseudoknots without compromising their efficiency. Some methods such as IPknot [9] and ProbKnot [10] use heuristics to predict also pseudoknotted structures.

#### Energy-based learning methods

The MFE structure for a sequence, given an energy model, is not necessarily the desired target structure. Energy models are not perfect because the thermodynamic parameters are calculated experimentally from many yet not sufficient number of samples. The ContraFold method [11] tries to learn new parameter sets and find the structure with respect to them, although the optimization is still with a dynamic programming algorithm.

#### Deep Learning methods

CDPFold [12] uses a convolutional neural network to predict a scoring matrix that is then fed to a dynamic programming algorithm to extract the dot-bracket structure. It can only predict non-pseudoknotted structures due to being limited to the dot-bracket notation, and also is not time-efficient for sequences longer than a few hundred bases because of the dynamic programming post-processing. Recently, E2Efold [13] proposed a deep neural network that outputs scores for all possible pairings in an RNA sequence and a differentiable post-processing network that converts the scores into a secondary structure. The score network of E2EFold had an architecture based on transformers [14] and convolutional layers. The post-processing tool was designed by convex relaxation of a discrete optimization problem to a continuous one. SPOT-RNA [15] is a deep learning model based on convolutional layers and custom 2D-BLSTM layers. This model considers also triplets (bases connected to two others) and non-canonical pairings. However it is limited to sequences shorter than 500 nucleotides (nt) due to the complexity of the model and memory limit. MXFold2 [16] is the most recent model which contains one-dimensional and two-dimensional convolutions and recurrent BiLSTM layers. The output has four different scores for each pair including helix stacking, helix opening, helix closing and unpaired region. The model that we propose in this paper is conceptually much simpler than the previous models, yet it results in very competitive performance.

#### Related problems

Learning-based methods dominate in related structure prediction problems. For example EternaBrain [17] uses reinforcement learning to address the RNA sequence design (inverse folding) problem. As another example, the recently proposed AlphaFold [18] set the new state of the art in predicting structures for proteins. The algorithm contains multiple deep learning components such as variational autoencoders, attention mechanism and convolutional networks.

### Problem definition

The problem of predicting the secondary structure of an RNA can be formulated as follows. Given a sequence of bases *q* = (*q*_1_, *q*_2_, … , *q*_*L*_), where each base *q*_*i*_ can take one of the four values A, U, C, G, the task is to predict a set of pairings {(*q*_*i*_, *q*_*j*_)} that define the secondary structure. For example, given a sequence CGUGUCAGGUCCGGAAGGAAGCAGCACUAAC, one needs to predict the pairings (*q*_2_, *q*_27_), (*q*_3_, *q*_26_), (*q*_4_, *q*_25_), (*q*_5_, *q*_24_), (*q*_10_, *q*_19_), (*q*_11_, *q*_18_), (*q*_12_, *q*_17_) which define the structure shown in Fig. 1.

**Fig 1.**
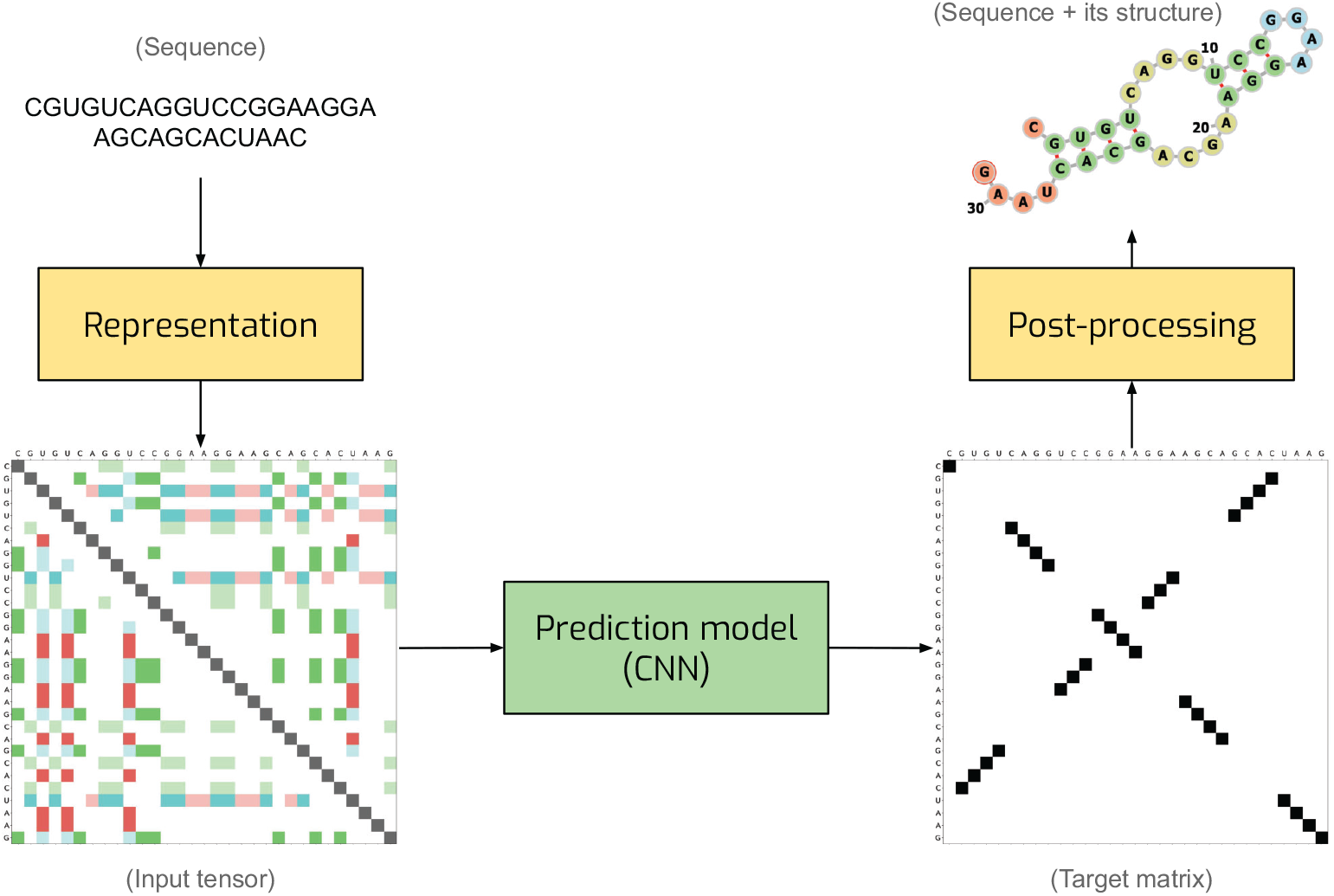
General illustration of our solution. We represent an RNA sequence as two-dimensional map with 8 channels with one-hot encoding. We process the map with a convolutional network which produces a score matrix for all possible pairings. Finally, we convert the score matrix into the RNA secondary structure.

There are a set of constraints that need to be satisfied:

- There are six possible types of pairings: (A, U), (U, A), (U, G), (G, U), (G, C), (C, G) (Watson-Crick and wobble pairing types).
- Each base can be either pairs with a single other base or unpaired. If base *i* is paired with base *j*, base *j* is paired with base *i*.
- The minimum distance for pairing is 3, that is |*i* − *j*| ≥ 3.

## Materials and methods

### Representing RNA sequences and secondary structure targets as tensors

The key component of our approach is the way we encode RNA sequences. We represent an RNA sequence *q* of length *L* as an *L* × *L* × 8 tensor *X* which can be viewed as a two-dimensional *L × L* map with eight channels (see the input tensor Fig 1). An eight-dimensional vector of features in location (*i, j*) of *X* is a one-hot representation of eight possible relations between bases *q*_*i*_ and *q*_*j*_ in positions *i* and *j*:

- Six channels indicate that base *q*_*i*_ can pair with base *q*_*j*_, that is pair (*q*_*i*_, *q*_*j*_) is one of the six possible combinations of bases (A, U), (U, A), (U, G), (G, U), (G, C), (C, G).
- One channel is used to indicate that *i* = *j*, i.e. this channel is set to ones only for the positions on the main diagonal of map *X*. The purpose of this channel is to ease detecting unpaired bases which we encode with non-zero elements on the diagonal of the target matrix.
- One channel indicates that a pairing between bases *i* and *j* is not possible due to a non-valid combination of bases, too short distance between two bases, or any other constraint.

We formulate the target for the model output as a binary *L* × *L* matrix *T* in which the *ij*-th element *t*_*ij*_ = 1 if bases *i* and *j* are paired and *t*_*ij*_ = 0 otherwise. *t*_*ii*_ = 1 means that base *i* is unpaired (see the target matrix Fig 1).

The advantage of the proposed representation is that it makes it equally easy to predict local and long-distance pairings. Local pairings are represented by non-zero elements in maps *X* and *Y* that are close to the main diagonal (see the target matrix in Fig. 1). Long-distance pairings correspond to locations in *X* and *Y* that are farther away from the main diagonal. Both types of structures can be easily detected by processing the input *X* with a convolutional neural network (CNN). CNN is also a powerful tool for detecting stem segments: blocks of consecutive bases paired with another block of bases. In our matrix representation, such pairings are represented by a sequence of non-zero elements in matrix *Y* which are either parallel or orthogonal to the main diagonal. These patterns can be easily detected with a CNN. Due to weight sharing, CNNs can process sequences of varying length and the processing is equivariant to translations of the input sequence. These are useful properties for our application.

### Prediction model

We represent each sequence as a 3-dimensional tensor (input tensor in Fig 1) which is the input of our prediction model. The output of the network is a 2-dimensional matrix (Target matrix in Fig 1) in which each element at (*i, j*) position shows the score for having (*i, j*) pairing in the predicted structure. Then, we extract the structure using our post-processing method. The

The prediction model takes an *L* × *L* × 8 tensor *X* as input and produces an output *Y* of shape *L* × *L*. The model starts with two convolution blocks followed by *M* residual blocks with skip connections and *N* residual blocks with shared weights (see Fig. 2). Each convolutional block is a convolutional layer with a 3 × 3 kernel and 32 output channels followed by batch-normalization and LeakyRelu activation function. To keep the size after each convolutional block unchanged, we have applied the required padding.

**Fig 2.**
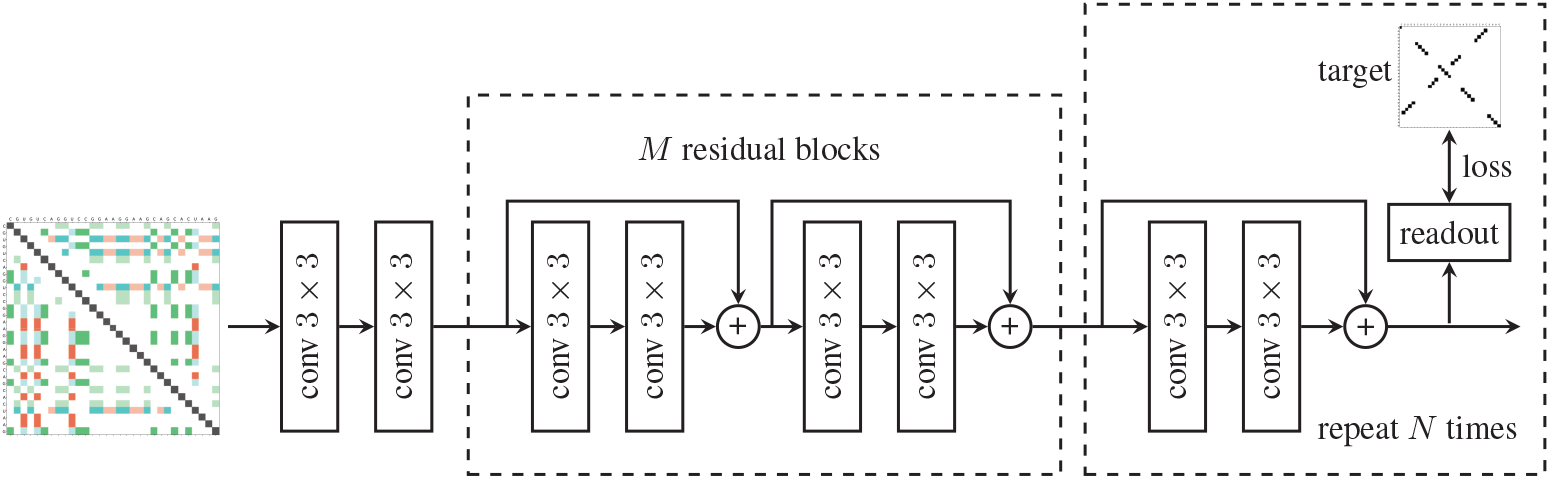
The architecture of the prediction model. “conv” denotes a convolutional layer followed by batch normalization and LeakyRelu nonlinearity. The readout layer is another conv block followed by a convolution layer both with kernel size 1. The loss is computed after each residual block with skip connections.

We encourage the network to arrive at the correct solution as fast as possible by computing the loss after each residual block. The model output *Y*_*n*_ after the *n*-th residual block is computed using a readout module which is a convolutional block followed by a convolution layer with one output channel. The loss function penalizes the difference between *Y*_*n*_ and the target matrix *T*. We use the mean-squared error as the loss:

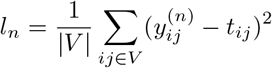

where 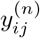 is the *ij*-th element of *Y*_*n*_ and *V* is a set of all pairs except for pairs with a non-valid combination of bases, too short distance between the bases or any other constraints. The mean-squared error was chosen because it produced the best results among other alternatives that we tried. The final loss is the average of the *N* intermediate loss values 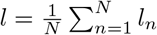.

### Post-processing

The output of the model is an *L* × *L* matrix which needs to be converted into a set of pairings that represent the secondary structure of an RNA sequence. We have used two alternative approaches for post-processing.

In the first approach, we try to extract a secondary structure in which each base is either paired to a single other base or unpaired (this condition holds for all RNA structures in the training datasets that we considered). The model output *Y* = *Y*_*N*_ is interpreted as a weighted adjacency matrix of a weighted graph in which the nodes correspond to bases and the weights of the edges reflect the chance that there is a pairing between the corresponding bases according to the model.^1^ Our goal is to find a set of edges without common nodes and with the option of including self-loops such that the sum of the weights is maximised. We solve this problem using the Blossom algorithm.^2^ This post-processing algorithm guarantees that each base is either paired with a single other base or unpaired. The problem with this post-processing is that it is computationally expensive, especially for long sequences.

In the second approach, we make a connection between base *i* and base *j* if the corresponding element *y*_*ij*_ of the model output has the maximum value in the *i*-th row of *Y*. This algorithm runs in time *O*(*L*^2^) but it often produces invalid structures because it does not guarantee the symmetry of pairings. This post-processing algorithm, however, yields similar results in terms of precision and recall compared to the Blossom post-processing (see Table 2). Therefore, we use it when we tune the hyperparameters of the model. We call this algorithm *Argmax post-processing*.

## Results

### Datasets

There are three commonly used datasets for RNA structure prediction.

1. RNAStralign [19] contains 37149 structures from eight RNA families with sequence lengths varying between 30 and 1851 nucleotides (nt). For sequences with multiple secondary structures, we randomly kept only one target secondary structure and therefore retained only 30451 samples. We split the dataset into 80% training, 10% validation, and 10% test sets (exactly as suggested in [13]) so that each RNA family had approximately the same representative fraction in each set as in the full dataset. Fig 3 shows the frequency of different lengths in this dataset in which the proportions are the same for all train, test, and validation sets. We use this dataset for training and testing purposes.
2. ArchiveII [20] contains 2975 samples with sequence lengths between 28 and 2968 nt from 10 RNA families (two additions to the RNAStrAlign families). We only tested our model on this dataset without any fine-tuning to evaluate it on a completely different sample set and compare with other methods.
3. BpRNA [15] contains 1305 samples shorter than 500 nt from several RNA families. We tested our model on this dataset without any fine-tuning to evaluate our model performance on samples from completely new families.

**Fig 3.**
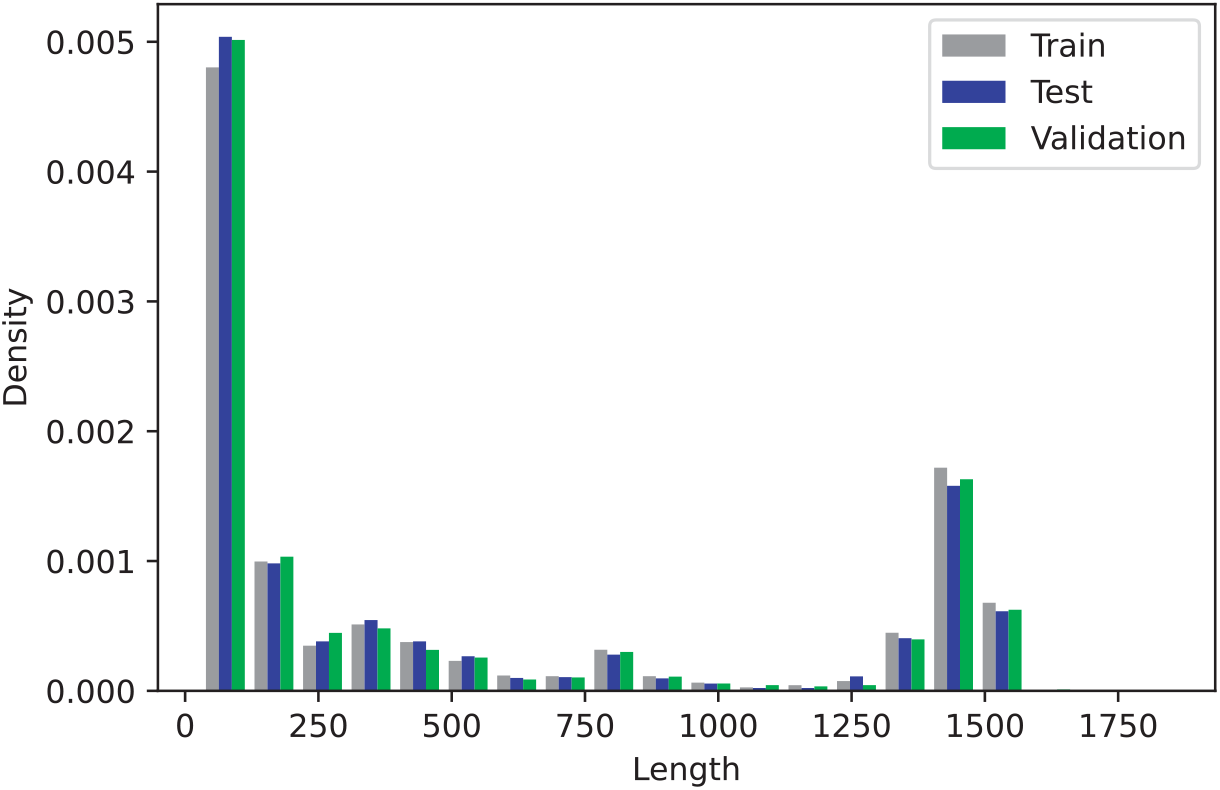
RNAStrAlign dataset lengths density for train, test, and validation sets. We use density instead of absolute number of samples because of different set sizes.

We evaluated the trained models using average precision, recall and F1-score, where precision reflects “how correct are our predicted pairings”, recall shows “how many of the target pairings our model could predict”, and F1-score is a harmonic average of the first two.

We trained the following variants of the proposed model.

1. CNNFold has *M* = 2 residual blocks and *N* = 2 shared residual blocks trained for 30 epochs on the whole trainset.
2. CNNFold-600 has *M* = 2 residual blocks and *N* = 2 shared residual blocks but trained for 400 epochs on the samples shorter than 600 nt.
3. CNNFold-600-big has *M* = 10 residual blocks and *N* = 2 shared residual blocks trained for 45 epochs on samples shorter than 600 nt. Due to the memory limit, we cannot use this model for long sequences.

We trained three different models using the Adam optimizer with learning rate 0.005. To avoid problems caused by the limited size of the GPU memory, we used mini-batches with varying sizes. We used only one sequence in a mini-batch for sequences longer than 1000 nt, while we used up to 16 samples in mini-batches containing shorter sequences. CNNFold and CNNFold-600-big have 95k and 317k parameters respectively while E2EFold and SPOT-RNA have 719k and 1746k parameters respectively.

While tuning the model, we found that CNNFold works slightly worse on short sequences (*L* ≤ 600) than CNNFold-600. CNNFold-600-big outperforms the other two models on short sequences. These results are presented in Table 1. Eventually, we use CNNFold-600-big to process sequences with *L* ≤ 600 and CNNFold to process sequences with *L* > 600. We call this ensemble of the two models CNNFold-mix.

**Table 1.** Results for our model trained on different parts of the RNAStrAlign training set.

| | F1, all data | F1, $L \leq 600$ | Weighted F1, all data |
| --- | --- | --- | --- |
| CNNFold-600-big | - | **0.970** | - |
| CNNFold-600 | 0.900 | 0.948 | 0.812 |
| CNNFold | **0.916** | 0.928 | 0.842 |
| CNNFold-mix | 0.936 | 0.970 | 0.863 |

The results on the RNAStrAlign dataset indicate that our model achieves significant improvements compared to the present state of the art (see Table 2). For example, the fraction of undetected pairings is only 0.093 for our model, which is less than two-fifths of the value 0.212 achieved by E2Efold. Our model achieves an impressive F1-score of 0.936 which is substantially higher than 0.821 of the previously best method on RNAStrAlign dataset. Fig 4 shows one randomly picked sample from 5S family. There are two other examples (S2 Fig and S3 Fig) in the supplementary materials. Not only are all the predictions visually close to their target structures, but they are also valid structures due to our post-processing method.

**Fig 4.**
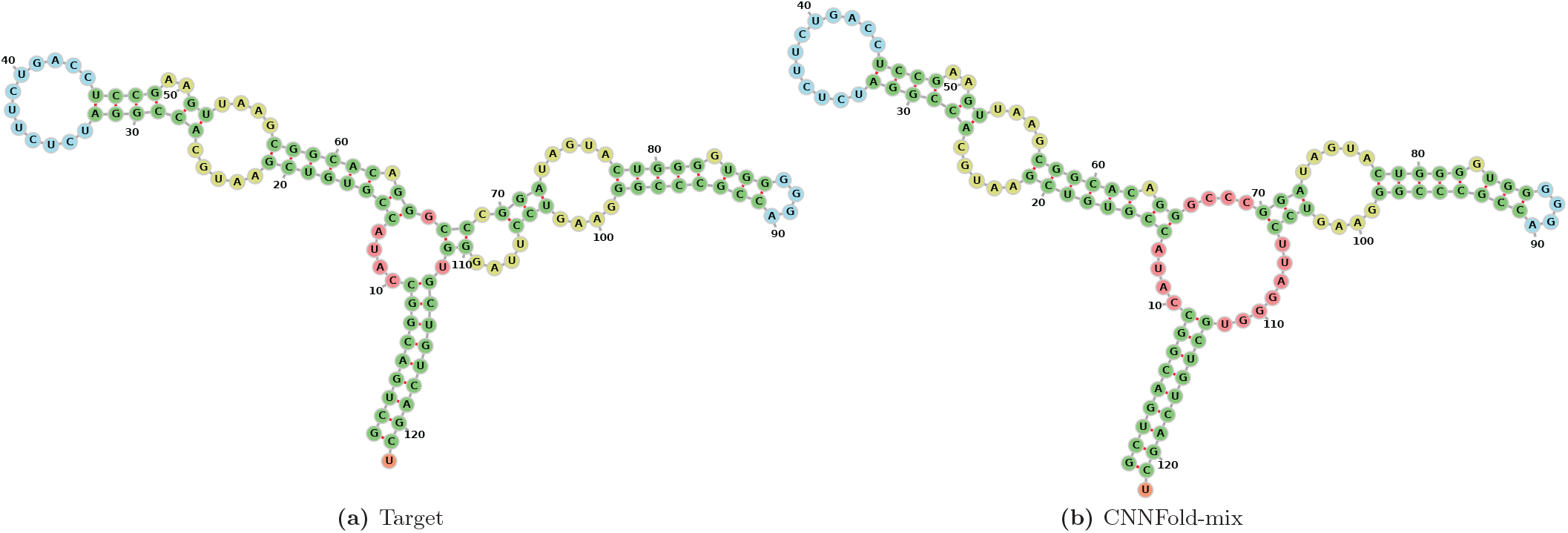
Visualization for E00001 from 5sRNA family. (a) is the target secondary structure and (b) is our prediction produced by CNNFold-mix with 93.4% accuracy. Structure diagrams are generated by the Forna [21] tool.

**Table 2.** Results on the RNAStrAlign dataset. “(S)” indicates the results when one-position shifts are allowed, that is for a base pair (*i, j*), the following predictions are also considered correct: (*i* + 1, *j*), (*i* − 1, *j*), (*i, j* + 1), (*i, j* − 1). The numbers for the comparison methods are from [13].

|  | Precision | Recall | F1 | Prec (S) | Recall (S) | F1 (S) | Weighted F1 |
| --- | --- | --- | --- | --- | --- | --- | --- |
| Mfold [5] | 0.450 | 0.398 | 0.420 | 0.463 | 0.409 | 0.433 | 0.366 |
| RNAfold [7] | 0.516 | 0.568 | 0.540 | 0.533 | 0.587 | 0.558 | 0.444 |
| RNAstructure [6] | 0.537 | 0.568 | 0.550 | 0.559 | 0.592 | 0.573 | 0.471 |
| LinearFold [22] | 0.620 | 0.606 | 0.609 | 0.635 | 0.622 | 0.624 | 0.509 |
| CDPfold [12] | 0.633 | 0.597 | 0.614 | 0.720 | 0.677 | 0.697 | 0.691 |
| CONTRAFold [11] | 0.608 | 0.663 | 0.633 | 0.624 | 0.681 | 0.650 | 0.542 |
| E2Efold [13] | 0.866 | 0.788 | 0.821 | 0.880 | 0.798 | 0.833 | 0.720 |
| CNNFold + Argmax | 0.955 | 0.861 | 0.900 | 0.955 | 0.872 | 0.902 | 0.812 |
| CNNFold-mix + Argmax | 0.956 | 0.912 | 0.932 | 0.958 | 0.915 | 0.934 | 0.863 |
| **CNNFold-mix + Blossom** | **0.975** | **0.907** | **0.936** | **0.978** | **0.909** | **0.938** | **0.872** |

Our model performs very well on both short and long sequences. One indication of this are the results presented in Table 1. To emphasize the performance on longer sequences, similarly to [13], we computed a weighted average of the F1-scores where the weight for a sequence of length *L*_*k*_ is *w*_*k*_ = *L*_*k*_/Σ_*k*_ *L*_*k*_. The weighted F1-score (see the last column of Table 2) indicates that our model works much better on long sequences compared to the previous methods. Fig. 5 shows a more in-detail scatter plot in which each point represents a sample with its length and F1-score.

**Fig 5.**
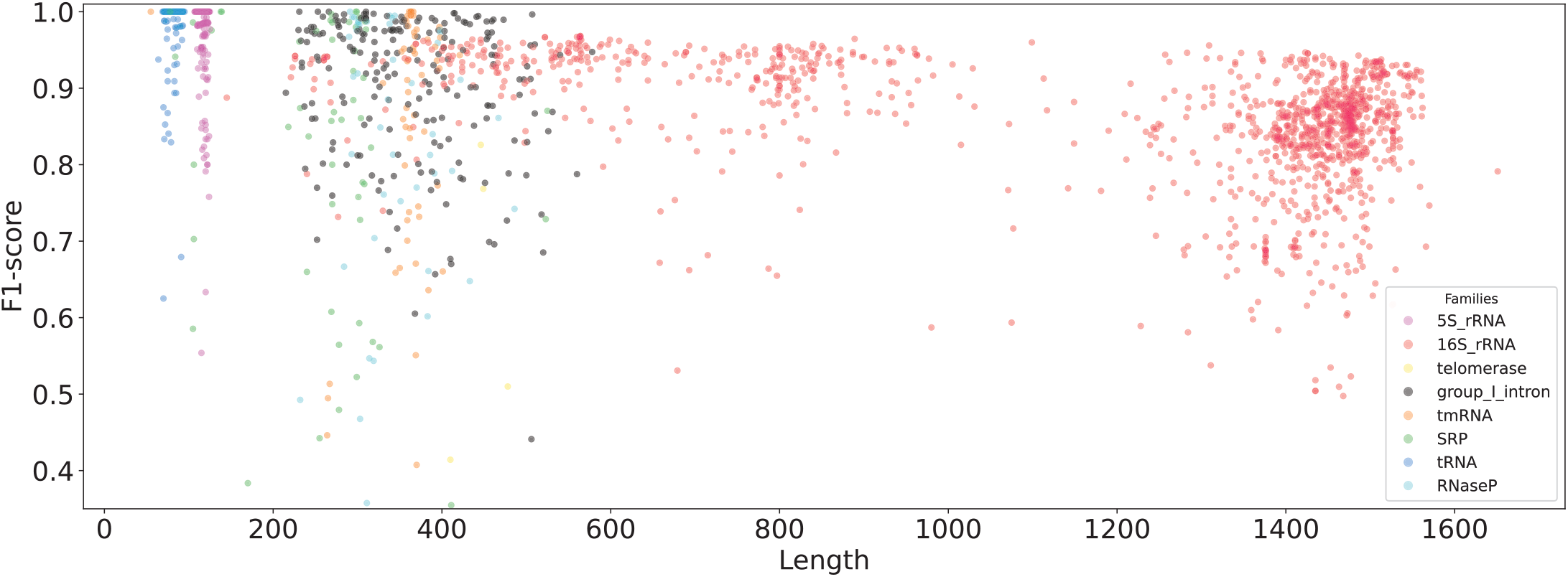
Scatter plot of the per-sequence F1-scores against the sequence lengths. Each point represents a sample and the model is CNNFold-mix. Colours indicate sequences from eight RNA families from RNAStrAlign.

To test how our model generalises to a dataset with a different distribution of sequences, we evaluated its performance on the popular ArchiveII dataset [20] (summarised in Table 3). The achieved F1-score is slightly smaller than the one on RNAStrAlign, but our model is clearly the best model among the competitors.

**Table 3.** Performance on the ArchiveII dataset.

|  | Precision | Recall | F1 |
| --- | --- | --- | --- |
| Mfold | 0.428 | 0.383 | 0.401 |
| CDPfold | 0.557 | 0.535 | 0.545 |
| RNAstructure | 0.563 | 0.615 | 0.585 |
| RNAfold | 0.565 | 0.627 | 0.592 |
| LinearFold | 0.641 | 0.617 | 0.621 |
| CONTRAFold | 0.607 | 0.679 | 0.638 |
| E2Efold | 0.734 | 0.660 | 0.686 |
| MXFold2 [16] | 0.790 | 0.815 | 0.800 |
| **CNNFold-mix** | **0.928** | **0.879** | **0.897** |

The SPOT-RNA group [15] did not report results on the ArchiveII dataset and replicating their model is not straightforward due to its complexity and the 500 nt length limit. However, we evaluated our model on the 1305 samples from the BpRNA dataset. without any retraining or fine-tuning. The achieved F1-score for CNNFold is 0.592 while the score reported in [15] was 0.630 after training on the same dataset.

Although our training dataset is different and our model is conceptually much simpler, the achieved F1-score is comparable with SPOT-RNA.

Our performance highly depends on the specific RNA families, with Telomerase and SRP being the hardest families for our model. In Fig. 5, we use different colours to show F1-scores for sequences from eight RNA families from the RNAStrAlign test set. The average F1-scores for different RNA families are shown in Table 4 for the RNAStrAlign and ArchiveII datasets.

**Table 4.** F1-scores obtained with CNNFold-mix for different RNA families.

| RNA Family | RNAStrAlign | ArchiveII |
| --- | --- | --- |
| 16S | 0.855 | 0.639 |
| 23S | – | 0.489 |
| 5S | 0.992 | 0.972 |
| Grp 1 Intron | 0.903 | 0.722 |
| Grp 2 Intron | – | 0.591 |
| RNaseP | 0.832 | 0.824 |
| SRP | 0.787 | 0.798 |
| telomerase | 0.615 | 0.755 |
| tmRNA | 0.830 | 0.871 |
| tRNA | 0.996 | 0.937 |

CNNFold-mix outpeforms other methods in predicting pseudoknotted structures. Out of the 3707 samples in our RNAStrAlign test set, 1413 are pseudoknotted, and we achieved an F1-score 0.857 on this subset. Although pseudoknotted structures are presumed to be more complex, CNNFold predicts them almost as well as non-pseudoknotted ones. A scatter plot of the pseudoknotted samples with respect to their lengths and their F1-score is presented in S1 Fig.

## Discussion

We have proposed a new learning-based method CNNFold for RNA secondary structure prediction independent of any energy model. Our results show that CNNFold significantly outperforms state-of-the-art methods E2Efold on RNAStrAlign and MXFold2 on ArchiveII datasets while achieving comparable results as the SPOT-RNA on the BpRNA without any fine-tuning on the dataset. Although the CNNFold model is less complex than the others (without any LSTM-like layer and with fewer parameters), it shows an outstanding performance thanks to the representation in which possible pairings are considered instead of only the sequence.

We believe that the method can be improved further to achieve an even better accuracy. One possibility is to take into account the length of the RNA sequence. This may have a positive impact on accuracy as the ensemble of models trained on different sequence lengths achieved the best performance in our experiments.

An important line of future research is to understand the limitations of the proposed method and other learning-based algorithms for RNA secondary structure prediction. The accuracy of the model is superb but it is important to understand how well the model can predict structural elements which are biologically important. For example, pseudoknots are difficult to predict, but missing any of them may have a significant effect on the functional properties of an RNA structure.

The ultimate goal of this line of research is to design new RNA sequences with the required functional properties. The results presented in this paper suggest that the proposed model can be a useful building block towards achieving this goal. It remains to be seen how well the model generalises to completely new RNA sequences that can be proposed in the design process. It may also be useful to extend the model to support multiple secondary structure predictions for a given sequence. This way one can increase the chance of finding RNA structures with the required functional properties.

## Acknowledgments

This work has been supported by Academy of Finland grant 311639, “Algorithmic Design for Biomolecular Nanotechnology (ALBION)”. The authors wish to acknowledge CSC – IT Center for Science, Finland, for computational resources.

---

**S1 Fig.**
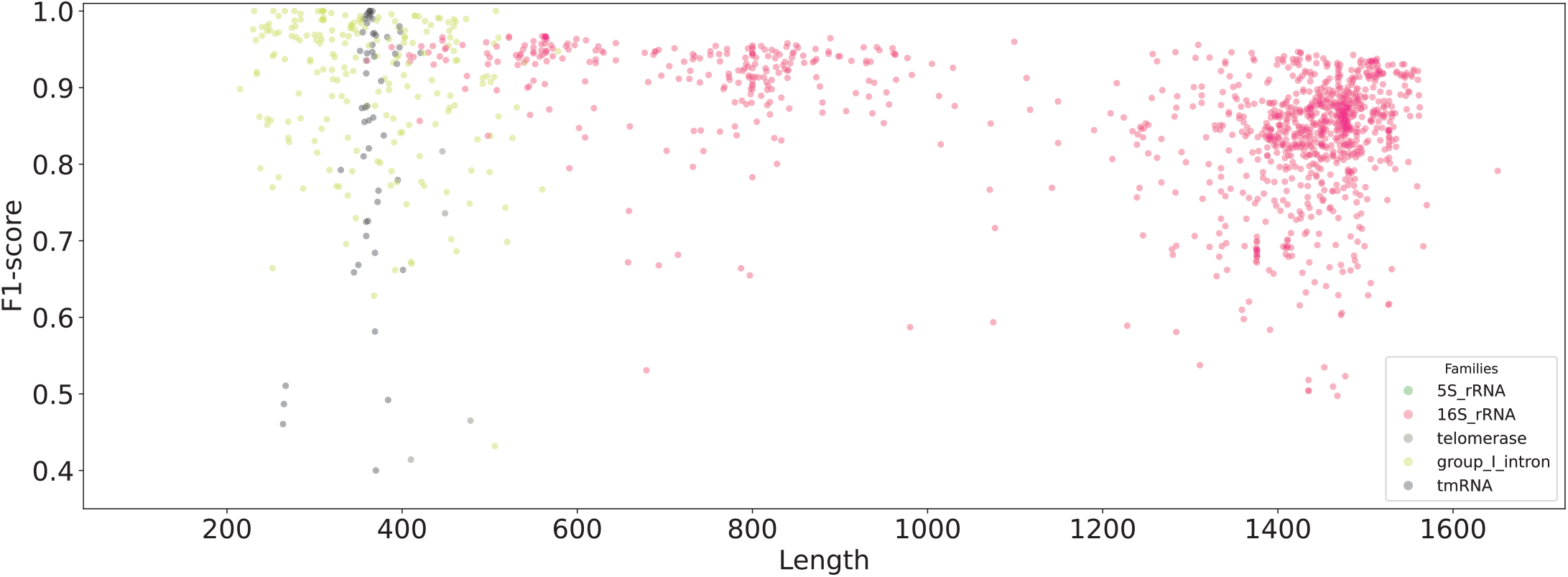
Scatter plot of the per-sequence F1-scores against the sequence lengths. Each point represents a sample and the model is CNNFold-mix. Colours indicate sequences from 6 RNA families from RNAStrAlign (only pseudoknotted structures).

**S2 Fig.**
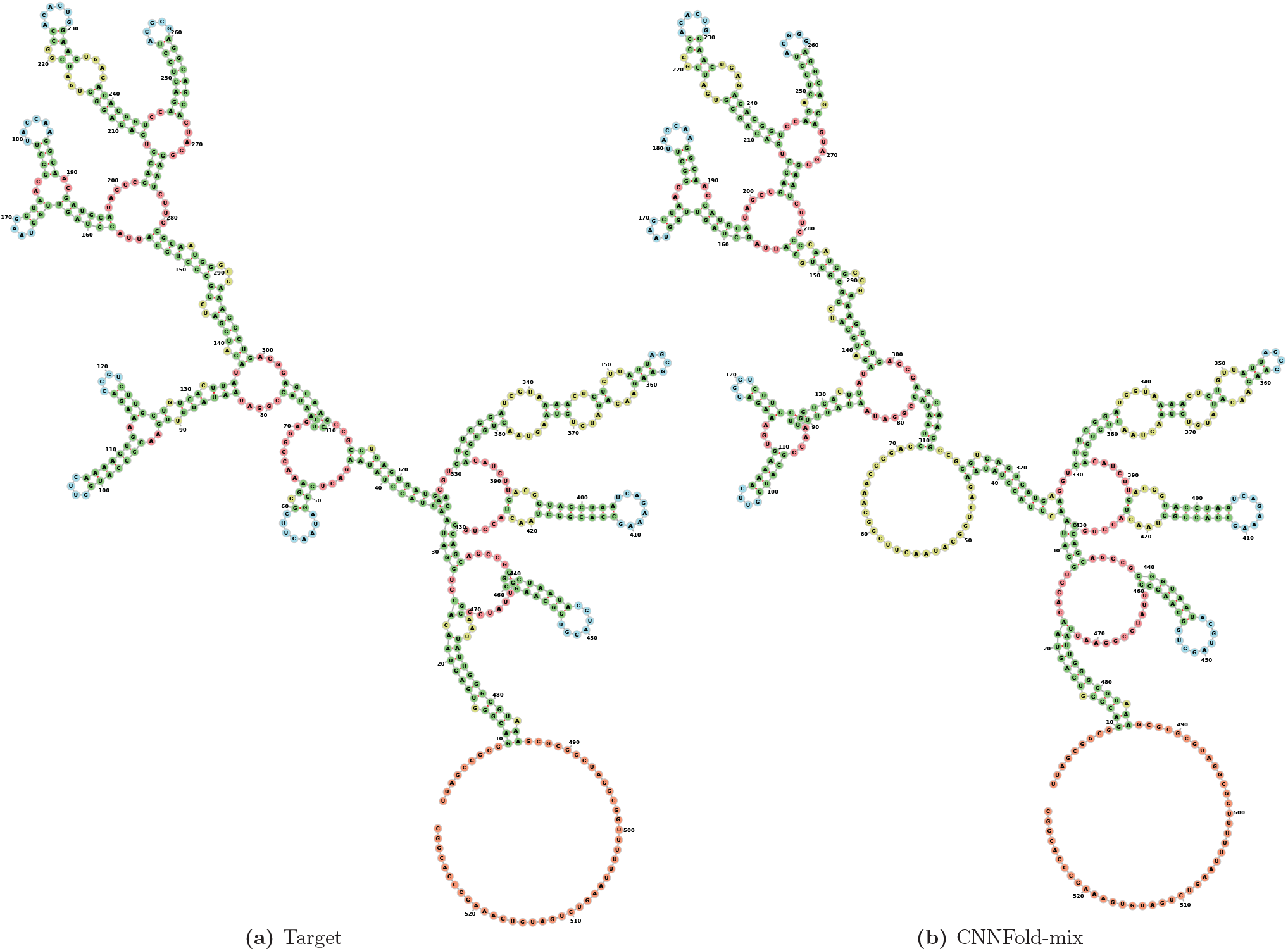
DQ923214 from 16sRNA family, accuracy of CNNFold-mix is 97.1% F1-score. (a) is the target structure and (b) is our prediction

**S3 Fig.**
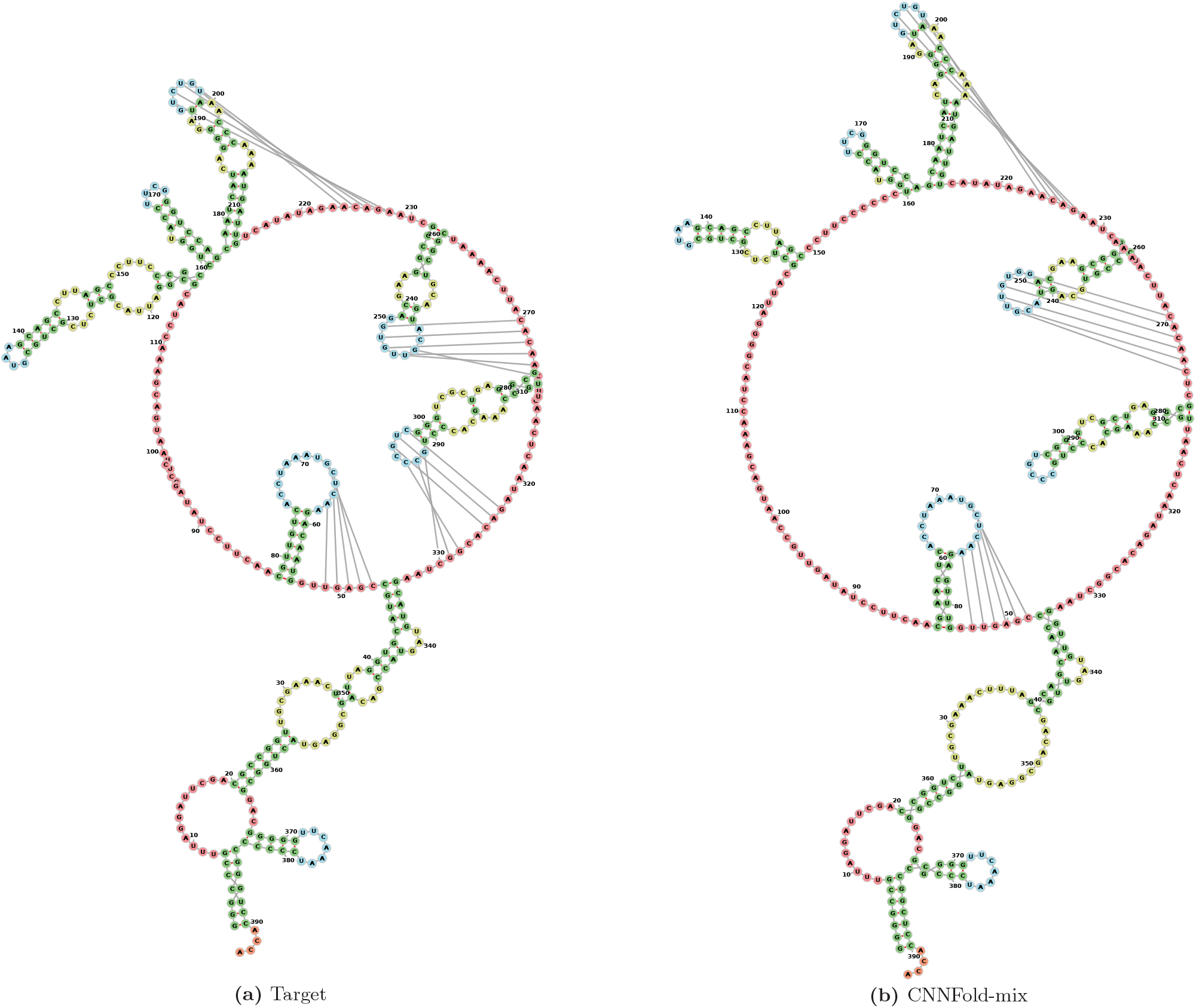
CP000076 with pseudoknots, F1-score of CNNFold-mix is 90.8%. (a) is the target structure and (b) is our prediction.

## Footnotes

1 To reduce the computational cost, we retain only *k* = 3 edges of maximum weight for each node.

2 We used available implementations of Blossom that do not support self-loops. To overcome this problem, we created a graph which contains two copies of the original weighted graph with the self-loops excluded and with additional connections between each pair of nodes representing the same node in the original graph. The weights of the additional connections are the weights of the corresponding self-loops multiplied by two.

